# The selective estrogen receptor modulator clomiphene inhibits sterol biosynthesis in *Arabidopsis thaliana*

**DOI:** 10.1101/2023.03.02.530820

**Authors:** Qing Wang, Kjell De Vriese, Sandrien Desmet, Jacob Pollier, Qing Lu, Alain Goossens, Danny Geelen, Eugenia Russinova, Geert Goeminne, Tom Beeckman, Steffen Vanneste

**Author notes:** These authors contributed equally (Qing Wang, Kjell De Vriese). **Author Contributions:** Q.W., K.D.V., Q.L. performed phenotyping analyses and experiments related to target identification. S.D. and J.P. performed and analyzed the GC-MS experiments. T.B., D.G., E.R., G.G. and S.V. conceptualized and supervised the experiments, and all authors contributed to the writing of the manuscript.

## Abstract

Sterols are produced via complex, multistep biosynthetic pathways involving similar enzymatic conversions in plants, animals and fungi, yielding a variety of sterol metabolites with slightly different chemical properties to exert diverse and specific functions. The role of plant sterols has been studied in the context of cell biological processes, signaling and overall plant development, mainly based on mutants. Due to their essential nature, genetic interference with their function causes pleiotropic developmental defects. An important alternative is to use a pharmacological approach. However, the current toolset for manipulating sterol biosynthesis in plants remains limited. Here, we probed a collection of inhibitors of mammalian cholesterol biosynthesis to identify new inhibitors of plant sterol biosynthesis. We provide evidence that imidazole-type fungicides, bifonazole, clotrimazole and econazole inhibit the obtusifoliol 14α-demethylase CYP51, that is highly conserved among eukaryotes. Surprisingly, we found that the selective estrogen receptor modulator, clomiphene, inhibits sterol biosynthesis, in part by inhibiting the plant-specific cyclopropyl-cycloisomerase CPI1. These results demonstrate that rescreening of the animal sterol biosynthesis pharmacology is an easy approach for identifying novel inhibitors of plant sterol biosynthesis. Such molecules can be used as entry points for the development of plant-specific inhibitors of sterol biosynthesis that can be used in agriculture.

## Introduction

Phytosterols mainly function as structural components in the plasma membrane where they regulate membrane permeability and fluidity, contribute to organizing the membrane in microdomains, and modulate the activity of membrane-bound enzymes (Cacas *et al*., 2012; Schaller, 2003; Simon-Plas *et al*., 2011).

In plants, many cellular processes are sterol-dependent, such as clathrin-mediated endocytosis (Konopka *et al*., 2008; Men *et al*., 2008)), cytokinesis (Boutte *et al*., 2010; Nakamoto *et al*., 2015), polarity (Men *et al*., 2008; Stanislas *et al*., 2015), and signaling (Simon-Plas *et al*., 2011). Consequently, phytosterols have been implicated in various developmental processes, such as embryonic and post-embryonic development (Clouse, 2000; Schaller, 2003), fertility (Azpiroz *et al*., 1998; Catterou *et al*., 2001), plant flowering (Schaller, 2003), growth (He *et al*., 2000; Schaller, 2003), seed germination (Guo *et al*., 1995), biotic- and abiotic stress responses (Han *et al*., 2009; Kumar *et al*., 2015; Pose *et al*., 2009; Senthil-Kumar *et al*., 2013), and the auxin-mediated regulation of cell polarity, gravitropism, endocytosis and auxin efflux (Men *et al*., 2008; Pan *et al*., 2009; Willemsen *et al*., 2003; Yang *et al*., 2013). In addition to its structural function, campesterol is also a metabolic precursor for the growth regulatory plant hormone brassinolide (Fujioka and Sakurai, 1997; Lindsey *et al*., 2003; Santner *et al*., 2009; Vriet *et al*., 2013).

The importance of phytosterols and brassinosteroids for plant growth and development is made clear by the severe phenotypes that are observed in mutants deficient in their biosynthesis. Early and late sterol biosynthesis mutants and BR-deficient mutants have been described, which all typically show severe dwarfism and defects in fertility, cell elongation, flowering and senescence. Additionally, Arabidopsis mutants that are defective in early phytosterol biosynthesis enzymes such as STEROL METHYLTRANSFERASE 1 (SMT1), CYP51G1, FACKEL (FK) and HYDRA1 (HYD1) are also deficient in embryogenesis and seed development, and cannot be rescued by BR treatment (Boutte and Grebe, 2009; Clouse, 2000; Diener *et al*., 2000; Souter *et al*., 2002). However, while mutants are a great asset for the study of phytosterol and BR biology in plants, they are not without drawbacks. For instance, the severe growth phenotypes that are typical for sterol- and BR-biosynthesis mutants make it difficult to dissect what is direct, and what is an indirect pleiotropic effect due to the often strong developmental phenotypes in the mutants. Therefore, an interesting alternative to study phytosterol and BR biology in plants is the use of small molecular inhibitors that target specific steps of their biosynthesis pathways.

Squalene is the common precursor for sterols in plants, animals and fungi. While the sterols found in the three eukaryotic kingdoms are highly diverse, their biosynthesis often involves similar metabolic steps. This similarity is illustrated by the ability of plant enzymes to complement yeast mutants in the corresponding enzyme (De Vriese *et al*., 2021; Diener *et al*., 2000; Kushiro *et al*., 2001), suggesting the distinct metabolic precursors can still dock the substrate binding pockets of these evolutionary distant enzymes. Consequently, sterol biosynthesis inhibitors that target specific sterol biosynthesis enzymes in fungi and mammals, often also inhibit sterol biosynthesis in plants. However, the molecular targets of these inhibitors are often much less defined, or their selectivity is relatively low due to divergence relative to their yeast and mammalian counterparts (De Vriese *et al*., 2021; He *et al*., 2003; Rozhon *et al*., 2013). For instance, reported oxidosqualene cyclase (OSC) inhibitors seem to non-selectively inhibit the activities of both cycloartenol synthase (CAS) and β-amyrin synthase (bAS) in plants (Ito *et al*., 2013). However, since the sterol biosynthesis pathways of plants, animals and yeast share many analogous conversion steps that are catalyzed by semi-conserved enzymes (Desmond and Gribaldo, 2009), several sterol biosynthesis inhibitors were found to be bioactive across kingdoms, albeit with distinct specificities. This is indeed the case for fenpropimorph, a morpholine-derived fungicide that is a known inhibitor of C-8,7 sterol isomerase (subnanomolar concentrations) and C-14 sterol reductase (micromolar concentrations) in yeast (Kerkenaar, 1990; Marcireau *et al*., 1990), which is also used as an inhibitor of FK, the plant C-14 sterol reductase, in plant research (He *et al*., 2003). However, fenpropimorph and similar morpholines only function at relatively high concentrations in plants (30 - 100 μM). Voriconazole, a triazole-type fungicide, inhibits members of the CYP51 superfamily of 14α-demethylase cytochrome P450 enzymes in both yeast (0.2 μM) (Saravolatz *et al*., 2003) and plants (1 μM) (Rozhon *et al*., 2013). Of these inhibitors, only fenpropimorph seems to be commonly used to manipulate sterol biosynthesis in plants.

Since the current library of characterized plant sterol biosynthesis inhibitors is rather limited (He *et al*., 2003; Rozhon *et al*., 2013), we set out to expand the catalog of plant sterol biosynthesis inhibitors. Therefore, we selected a subset of compounds that target different steps of the mammalian cholesterol biosynthesis pathway that were recently identified in a screen in a human cell system (Korade *et al*., 2016). We provide the proof-of-concept for several imidazoles as putative inhibitors of CYP51. Moreover, we identified the Selective Estrogen Receptor Modulator clomiphene as a novel inhibitor plant sterol biosynthesis in part by inhibiting the plant-specific cyclopropyl-cycloisomerase CPI1. These findings illustrate the principle that the wide array of, often commercially available, animal sterol biosynthesis inhibitors can be exploited for the efficient identification of novel plant sterol biosynthesis inhibitors

## Results

Chemical screens in human cell systems resulted in an expanded list of inhibitors that target different steps of cholesterol biosynthesis (Kim *et al*., 2016; Korade *et al*., 2016). The enzymatic steps of cholesterol biosynthesis in animals generally strongly resemble those of the plant phytosterol biosynthesis pathway (Desmond and Gribaldo, 2009). Therefore, we postulated that inhibitors of human cholesterol biosynthesis and/or fungal lanosterol biosynthesis could target analogous steps in phytosterol biosynthesis. We selected several representative compounds that target distinct cholesterol/lanosterol biosynthesis enzymes in humans and yeast (Kim *et al*., 2016; Korade *et al*., 2016), to evaluate their potential as inhibitors of plant sterol biosynthesis (Table 1). As positive controls we included several imidazoles, a class of molecules that is rich in inhibitors of a variety of Cytochrome P450s, such as CYP51.

**Table 1.** List of putative sterol biosynthesis inhibitors selected for analysis in Arabidopsis.

| Compound | Other bioactivities | Mammalian target in sterol biosynthesis | Analogous Arabidopsis enzyme(s) |
| --- | --- | --- | --- |
| Bifonazole | imidazole antifungal | CYP51A1 | CYP51 |
| Clotrimazole | imidazole antifungal | CYP51A1 | CYP51 |
| Econazole | imidazole antifungal | CYP51A1 | CYP51 |
| Clomiphene | Selective Estrogen Receptor Modulator (SERM) | C-8,7 isomerase/DHCR24 | HYD1/DWF1 |
| Tamoxifen | SERM | C-8,7 isomerase/DHCR24 | HYD1/DWF1 |
| Perphenazine | antipsychotic, CaM antagonist | C-8,7 isomerase/DHCR24 | HYD1/DWF1 |
| Fluphenazine | antipsychotic, CaM antagonist | C-8,7 isomerase | HYD1 |
| Acitretin | retinoid antipsoratic | DHCR24 | DWF1 |
| Doxepin | antipsychotic | DHCR24 | DWF1 |
| Trifluoperazine (TFP) | antipsychotic, CaM antagonist | DHCR24 | DWF1 |

### Hypocotyl and root phenotypes highlight putative phytosterol biosynthesis inhibitors among a set of cholesterol biosynthesis inhibitors

As an indirect readout of defective phytosterol biosynthesis (Rozhon *et al*., 2013), we monitored the ability of the inhibitors to reduce hypocotyl length of etiolated Arabidopsis seedlings (Fig. 1A-B; Supplementary Fig. S1A). Bifonazole, clotrimazole and econazole strongly reduced hypocotyl length and were already effective at 0.5 µM concentrations (Fig. 1A). Clomiphene had an effect at higher concentrations (5 – 10 µM range) and tamoxifen reduced hypocotyl length only at 10 µM (Fig. 1B). Another putative C-8,7 isomerase/DHCR24 inhibitor, Perphenazine, as well as the DHCR24 inhibitors fluphenazine, acitretin, trifluoperazine and doxepin had no significant effects on hypocotyl elongation, even at the highest concentration (10 µM) tested (Fig. 1B and Supplementary Fig. S1A), suggesting that the latter are not effective inhibitors of phytosterol biosynthesis.

**Figure 1.**
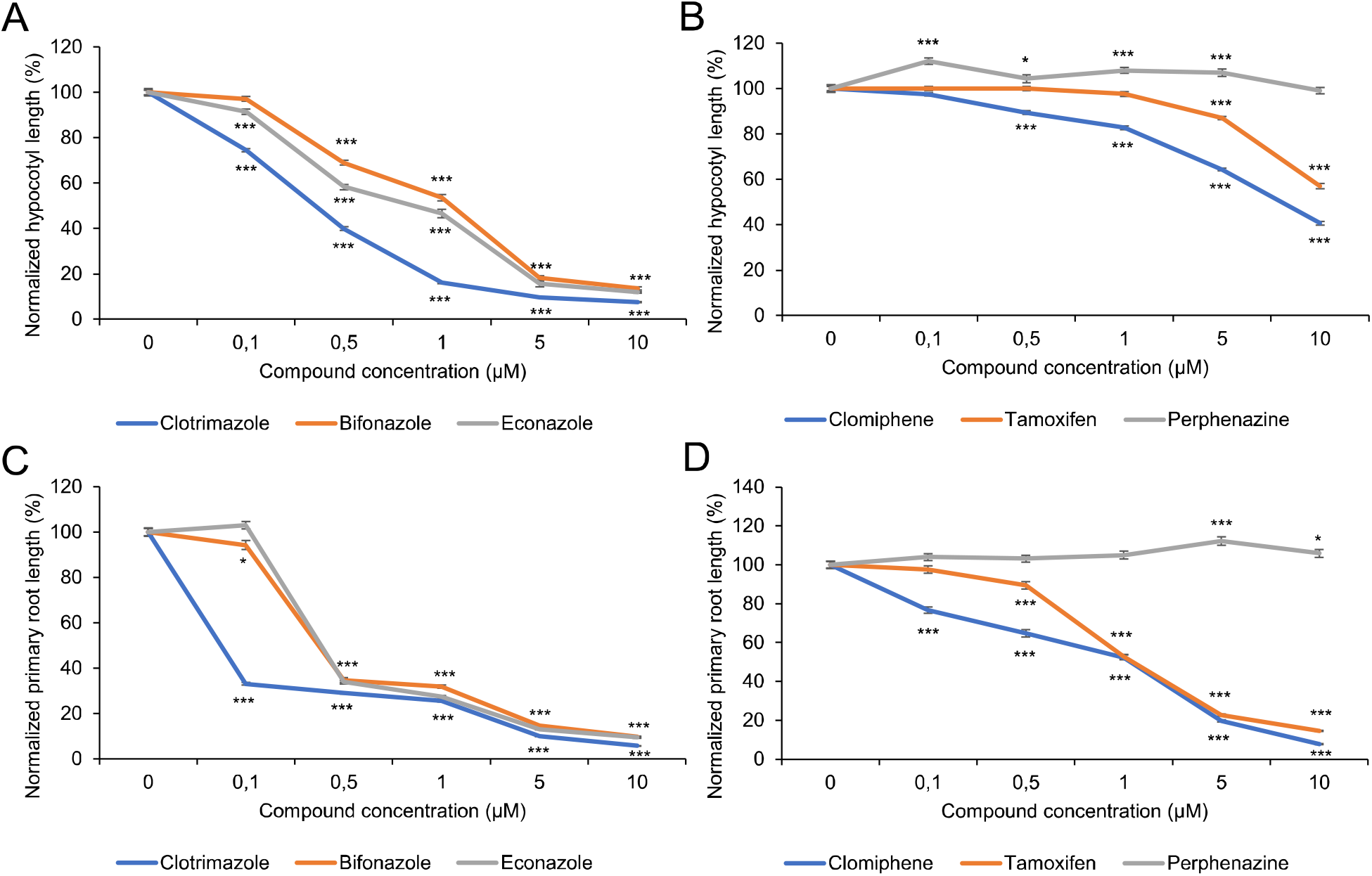
Effect of putative sterol biosynthesis inhibitors on hypocotyl and root length. **(A-B)** Dose-response curves of hypocotyl lengths of seedlings treated with (A) Clotrimazole, Bifonazole or Econazole and (B) Clomiphene, Tamoxifen or Perphenazine. Seedlings were grown for 8 days in the dark on ½ MS medium supplemented with the respective inhibitors at indicated concentrations. **(C-D)** Dose-response curves of primary root lengths of wild-type seedlings (Col-0) treated with and (C) Clotrimazole, Bifonazole or Econazole and (D) Clomiphene, Tamoxifen or Perphenazine. Wild type seedlings (Col-0) were grown for 7 days under continuous illumination on ½ MS medium supplemented with the respective inhibitors at indicated concentrations. Averages for each condition are depicted (n = 33 – 56) relative to the DMSO control (0.1 %). Error bars indicate ±SEM. Student’s t-test p-values: *p < 0.05, **p < 0.01, ***p < 0.001.

Next, we analyzed the effects of these inhibitors on root growth of light grown seedlings (Fig. 1C-D; Supp. Fig. S1B). In comparison to seedlings grown in the presence of 0.1% DMSO, several of the selected compounds strongly inhibited root growth. At concentrations as low as 0.1 µM, clotrimazole reduced the primary root length by more than 50% compared to the control, while, bifonazole and econazole did so at 0.5 µM (Fig. 1C). Clomiphene and tamoxifen also strongly reduced the primary root length, albeit at higher concentrations (1 µM and above) than the imidazoles (Fig. 1D). At concentrations of 5 µM and higher, clotrimazole, clomiphene and tamoxifen caused severe reductions in primary root length. Consistently with the low bioactivity in the hypocotyl elongation assay, acitretin, doxepin, perphenazine, fluphenazine and trifluoperazine had no obvious effect on the primary root length at the highest tested concentration of 10 µM (Supplementary Fig. S1B). Due to the low bioactivity in our assays, we did not further pursue acitretin, doxepin, perphenazine, fluphenazine and trifluoperazine in the subsequent analyses.

### Inhibitors that interfere with root growth cause cell division orientation defects in the root meristem

The defective sterol biosynthesis is often associated with defective cell division orientation, such as is the case for *fk* and *smt2smt3* double mutant (Jang *et al*., 2000; Pullen *et al*., 2010; Souter *et al*., 2002). Therefore, we visualized the root meristem organization of inhibitor treated roots using ABCB19-GFP as a plasma membrane marker (Fig. 2A-F). We focused on the inhibitors that caused reductions in the growth assays (Fig. 1C,D). The typical regular organization of the meristem was disrupted by all inhibitors, as indicated by the appearance of aberrant cell division orientations (Fig. 2A-F).

**Figure 2.**
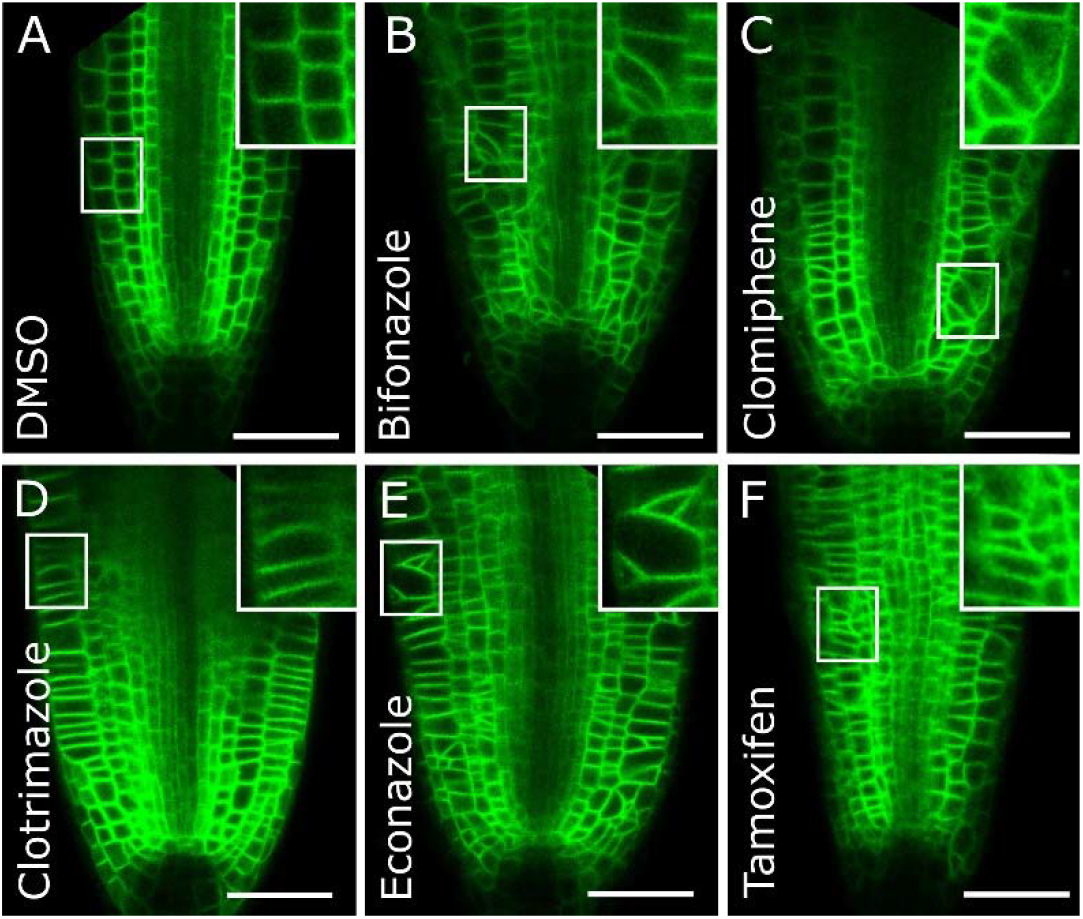
Perturbed cell division orientation in the root meristem after inhibitor treatment. **(A-F)** Root organization of 5 day-old ABCB19-GFP seedlings grown on 0.5 x MS supplemented with (A) DMSO (0.1%), (B) Bifonazole (1 µM), (C) Clomiphene (1 µM), (D) Clotrimazole (1 µM), (E) Econazole (1 µM), and (F) Tamoxifen (1 µM). Scale bar = 50 µm.

While bifonazole, clomiphene, econazole, and tamoxifen treatment potently disrupted cell division orientations in the root meristem, this was less obvious upon clotrimazole treatment (Fig. 2D). These cell division orientation defects are reminiscent of mutants defective in sterol biosynthesis (Jang *et al*., 2000; Pullen *et al*., 2010; Souter *et al*., 2002).

### GC-MS analysis reveals disturbed sterol composition in Arabidopsis seedlings after inhibitor treatment

Next, we determined the impact of the compounds on the sterol composition of Arabidopsis seedlings (Fig. 3). Seedlings were transferred for 5 days to liquid medium containing 0.5 µM and 5 µM of bifonazole, clomiphene, clotrimazole and econazole. Sterols were extracted and analyzed via gas chromatography-mass spectrometry (GC-MS). The major peaks in the GC-MS chromatograms corresponded to the three major phytosterols in Arabidopsis (campesterol, stigmasterol, and β-sitosterol), and the sterol biosynthesis intermediate isofucosterol (Fig. 3A). β-amyrin, which was added to the samples as an internal standard, and α-tocopherol (vitamin E) eluted in the same range as the major plant sterols (Fig. 3A). The identification of these metabolites was based on a NIST database search with their EI-MS spectra (Supplementary Fig. S2).

**Figure 3.**
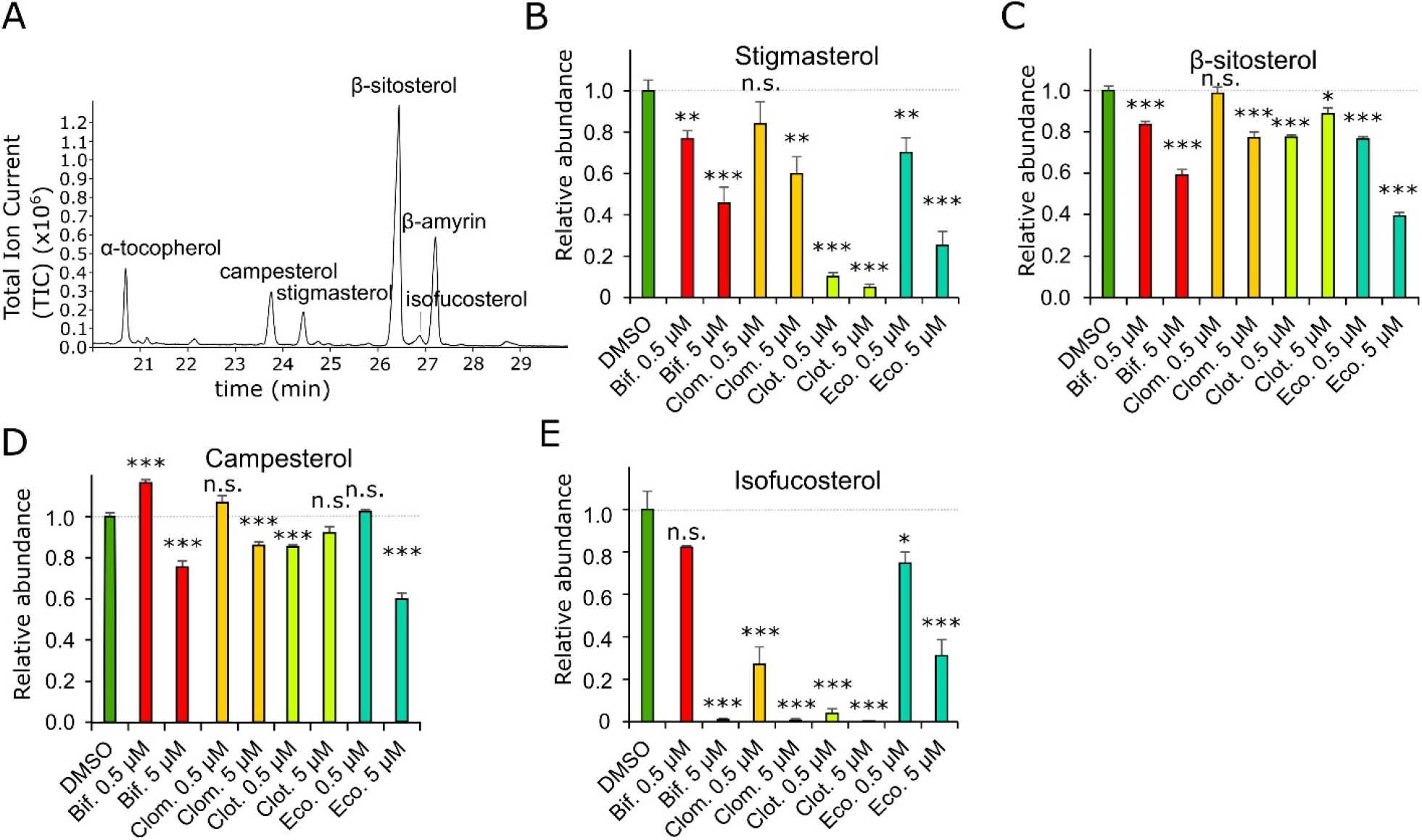
Relative quantification of the major sterols after inhibitor treatment. **(A)** Representative Total Ion Current (TIC) GC-MS chromatogram of a DMSO-treated WT Arabidopsis seedlings sample, with indication of α-tocopherol, campesterol, stigmasterol, β-sitosterol and β-amyrin. **(B-E)** Peak intensities of **(B)** stigmasterol, **(C)** β-sitosterol, **(D)** campesterol and **(E)** isofucosterol in WT seedlings treated with two concentrations (0.5 or 5 µM) of Bifonazole (Bif.), Clomiphene (Clom.), Clotrimazole (Clot.) and Econazole (Eco.), relative to the peak intensities in the DMSO control. Error bars represent ±SEM, *n* = 5. Student’s t-test p-values: *p < 0.05, **p < 0.01, ***p < 0.001. Non-significant (n.s.) p> 0.05.

From these analyses, it was clear that most of the tested compounds had at least some effect on the sterol composition of these three major phytosterols, and isofucosterol (Fig. 3B-E). The largest sterol disturbances were found in the stigmasterol levels, for which generally strong reductions were observed in samples treated with the imidazoles, with more modest effects for clomiphene (Fig. 3B). The relative levels of β-sitosterol, the precursor of stigmasterol, were generally less affected than those of stigmasterol (Fig. 3C). The effect of the inhibitors on campesterol levels was mostly modest, and even slight induction was noted for the 0.5 μM bifonazole treatment (Fig. 3C,D). All compounds strongly reduced the levels of the β-sitosterol-precursor isofucosterol (Fig. 3E). These data indicate that clomiphene and all tested imidazoles have an impact on sterol biosynthesis, either directly, or indirectly.

### Clotrimazole, bifonazole and econazole are inhibitors of CYP51 activity in Arabidopsis

To infer the sterol biosynthesis step that is most likely targeted by these inhibitors, we revisited the TIC GC-MS chromatograms for each treatment, in search of new peaks that likely correspond to sterol biosynthesis intermediates, and derivatives thereof. In the TIC GC-MS chromatograms of all imidazole-treated samples, we observed the appearance of up to 5 new peaks (Fig. 4A; Supplementary Fig. S3A,B).

**Figure 4.**
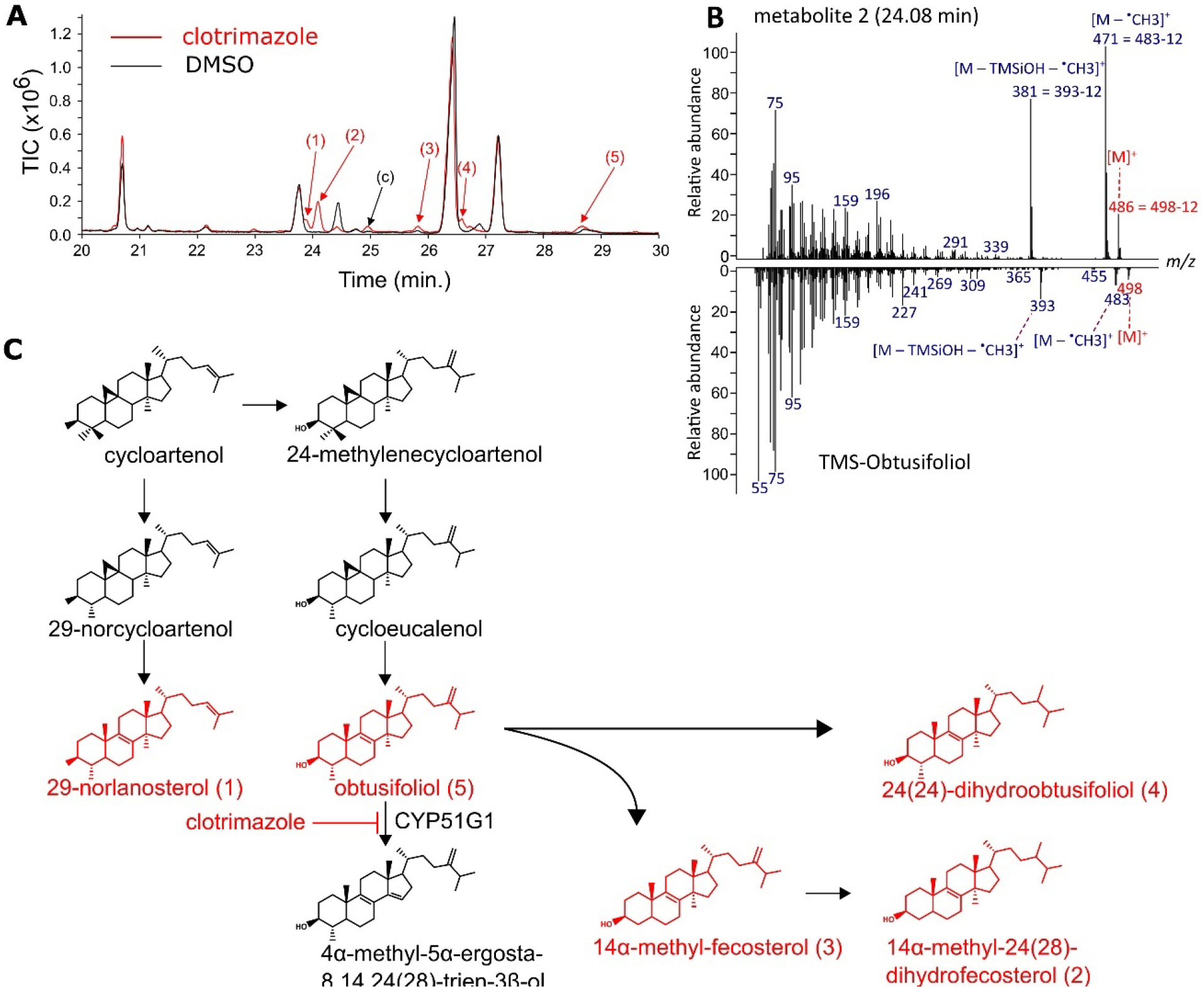
Clotrimazole target identification. **(A)** Overlay of a representative TIC GC-MS chromatogram of a control sample (black) and samples treated with 5 μM clotrimazole (red). Five new peaks induced by clotrimazole treatment are indicated by numbers for which the EI-MS spectra are presented in (B) and Supplementary Fig. 4, and are referred to as “metabolite 1” to “metabolite 5”. The peak indicated by (c) is a contaminant. **(B)** Comparison of the EI-MS profile of metabolite 2 to TMS-Obtusifoliol. Reference EI-MS profile was obtained from NIST ’20. **(C)** Model explaining how inhibition of CYP51G1 by clotrimazole treatment could lead to accumulation of the metabolites 1 tos 5.

The EI-MS profiles of most of these metabolites displayed prominent ions at [M-^•^CH3]^+^ and [M-TMSiOH-^•^CH3]^+^ (Fig. 4B; Supplementary Fig. S4A-D), which is indicative of Δ^8^ sterols with a 14α-methyl group (Goad and Akihisha, 1997). The accumulation of Δ^8^ sterols with a 14α-methyl group is consistent with clotrimazole, bifonazole and econazole inhibiting 14α-demethylase activity in Arabidopsis (AtCYP51G1), a deeply conserved target of many azoles (Crowley and Gallagher, 2014; Lamb *et al*., 2001). A NIST ’20 database search revealed that the EI-MS spectrum of metabolite 5 matched for 59.98% with that of TMS-obtusifoliol (Supplementary Fig. S4A), the preferred target of AtCYP51G1.

Metabolite 2 was the major accumulating metabolite not only for clotrimazole, but also for bifonazole and econazole treatments (Fig. 4A, Supplementary Fig. S3A-B) and had an EI-MS spectrum that was also highly similar to that of TMS-obtusifoliol (NIST20) (Fig. 4B). The major differences between both spectra were at the level of its molecular ion [M]^+^, and the fragments [M – ^•^CH3]^+^ and [M – TMSiOH – ^•^CH3]^+^, all of which had an *m/z* value that is 12 Da lower than the corresponding ions in the TMS-obtusifoliol spectrum, suggesting that it contains one carbon atom less than obtusifoliol, yielding C_29_H_50_O. This chemical formula corresponds to that of 14α-methyl-24(28)-dihydrofecosterol, the major metabolite accumulating in *atcyp51g1* mutants (Kim *et al*., 2005).

The biosynthesis of 14α-methyl-24(28)-dihydrofecosterol from obtusifoliol requires C4α-demethylation and C24(28) double bond reduction (Fig. 4C). The C4α-demethylation of obtusifoliol is also seen in the *Atcyp51g1* mutant with the accumulation 14α-methyl-fecosterol (C_29_H_48_O) (Kim *et al*., 2005). The molecular ion of both metabolite 1 and 3, i.e., *m/z* 484 (Supplementary Fig. S4B,C), corresponds with a chemical formula of C_29_H_48_O and a double bond equivalent (DBE) of 6, which indicates the presence of 2 double bonds. Both metabolites could thus be 14α-methyl-fecosterol. However, the prominent peak at *m/z* 69 in metabolite 1 (Supplementary Fig. S4B) is indicative for the presence of a Δ^24(25)^-double bond in the side chain of 14α-methylsterols (Goad and Akihisha, 1997), suggesting that metabolite 1 is 29-norlanosterol. By deduction, metabolite 3 then likely corresponds to 14α-methyl-fecosterol.

Metabolite 4 displayed a molecular ion at *m*/*z* 500 (Supplementary Fig. S4D), matching a chemical formula of C_30_H_52_O and a DBE of 5, indicating a single double bond. This matches the main features of 24(28)-dihydro-obtusifoliol, another metabolite that also accumulates in the *cyp51* mutant (Kim *et al*., 2005).

The accumulation of 14α-methyl sterols is a tell-tale sign of inhibition of CYP51 activity (Kim *et al*., 2005; Maillot-Vernier *et al*., 1990; Taton *et al*., 1988). Therefore, it can be concluded that bifonazole, clotrimazole, and econazole inhibit the Arabidopsis CYP51 orthologue (CYP51G1). That clotrimazole accumulated additional 14α-methyl sterols probably reflects its higher bioactivity compared to bifonazole, and econazole.

### Clomiphene inhibits CPI activity in Arabidopsis

In contrast to the imidazole TIC GC-MS chromatograms, no 14α-methyl-24(28)-dihydrofecosterol (2) peak appeared in the clomiphene samples. Instead, five induced peaks appeared in the clomiphene-treated samples that were not induced in the imidazole-treated samples (Fig. 5A; pink arrows), suggesting that clomiphene has a different target than the imidazoles. The EI-MS spectrum of the largest of these peaks **(10)** was a mixed spectrum of two metabolites (metabolites 10a and 10b). After deconvolution, we found via database searches that the fragment ions for these metabolites corresponded to the fragmentation spectra of TMS-cycloeucalenol and TMS-α-amyrin (Fig. 5B,C).

**Figure 5.**
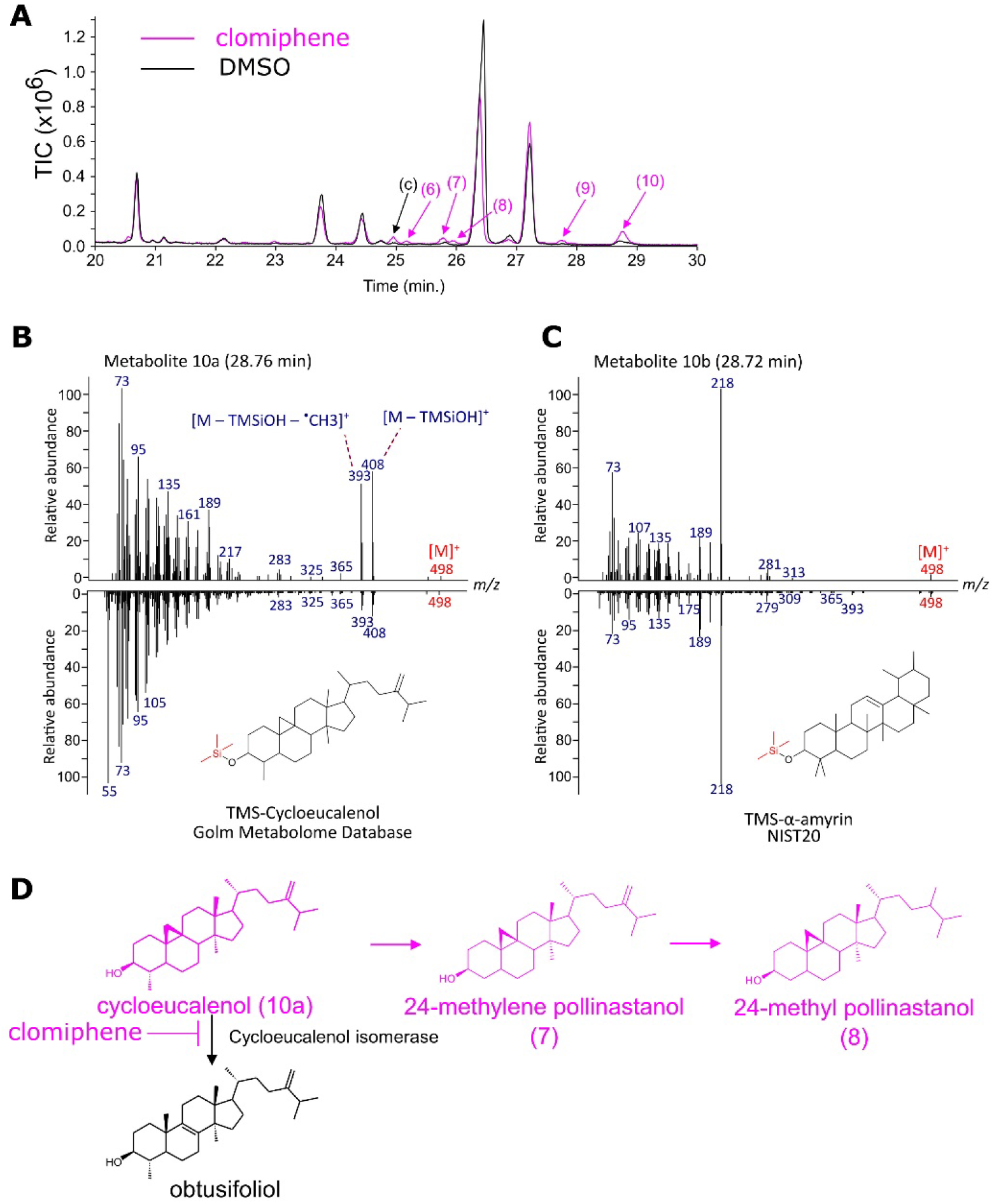
GC-MS based identification of CPI as a clomiphene target. (**A**) Overlay of a representative TIC GC-MS chromatogram of the control sample (black) and the samples treated with 5 μM (pink) clomiphene. Pink arrows indicate the positions of the accumulating metabolites. New peaks, induced by inhibitor treatment are indicated by numbers for which the EI-MS spectra are presented here and in **Supplementary Fig. S5**. (c) = contaminant. (**B**,**C**) Comparison of the EI-MS profile of Metabolite 10a to TMS-Cycloeucalenol (**B**) and Metabolite 10b to TMS-α-amyrin (**C**). Reference EI-MS profiles were obtained from the Golm Metabolome Database and NIST20. (**D**) Model explaining how clomiphene treatment could lead to accumulation of cycloeucalenol, 24-methylene pollinastanol and 24-methyl pollinastanol.

Similarly to TMS-cycloeucalenol, metabolite 7 and metabolite 8 showed two intense [M-TMSiOH]^+^ and [M-TMSiOH-^•^CH_3_]^+^ fragment ions (Supplementary Fig. S5A,B), indicating these metabolites are also 9β,19-cyclopropanesterols (Goad and Akihisha, 1997), related to cycloeucalenol. Based on the *m/z* of their molecular ions, one is likely its demethylated version, 24-methylene pollinastanol (metabolite 7) and the other differing from the previous by a reduced double bond, 24-methyl pollinastanol (metabolite 8). The identity of these metabolites was further supported by a shared fragment ion at *m/z* 269, representing the sterol backbone ([M-TMSiOH-SC]^+^) and a fragment ion unique to 24-methyl pollinastanol at *m/z* 220 (Böhme *et al*., 1997). Jointly, the accumulation of cycloeucalenol, and its aberrant metabolites 24-methylene pollinastanol and 24-methyl pollinastanol, suggests that clomiphene inhibits cycloeucalenol cycloisomerase (CPI) activity (Fig. 5D).

The spectrum of metabolite 6 was very similar to that of 24-methylene lophenol (C_29_H_48_O) (Zu *et al*., 2021), including major fragment ions (*m/z* 343, 255 and 229) (Supplementary Fig. S5C).This suggests that the structure of metabolite 6 is closely related to 24-methylene lophenol. The identity of metabolite 9 (Supplementary Fig. S5D) could not be reliably resolved due to low abundance. The accumulation of metabolites that are unrelated to cycloeucalenol indicates that clomiphene targets multiple steps in the phytosterol biosynthetic pathway.

## Discussion

Phytosterol biosynthesis is orchestrated by complex, branched pathways that are regulated by a wide range of enzymes, which makes the study of these processes often difficult. While several mutants are available that are defective in specific steps of phytosterol biosynthesis, these mutants often show severe growth phenotypes and are thus not ideal to study sterol biosynthesis defects later in a plant’s life. Unfortunately, the current toolset of sterol biosynthesis inhibitors in plants is limited (De Vriese *et al*., 2021). The close homology of the structures and the relative conservation of enzymes involved, explains why imidazoles that are known to block sterol biosynthesis in animals and/or fungi can also interfere with sterol biosynthesis in plants (He *et al*., 2003; Rozhon *et al*., 2013). While differences in structure and divergence of the catalytic centers may cause shifts in specificity, inhibitors of human sterol biosynthesis should thus be enriched in molecules that can interfere with plant sterol biosynthesis.

We exploited this principle by exploring a set of putative mammalian cholesterol biosynthesis inhibitors and identified several new inhibitors of plant sterol biosynthesis. Based on morphological and biochemical effects we demonstrate that the imidazoles, bifonazole, clotrimazole, and econazole, and the selective estrogen receptor modulator, clomiphene, are potent inhibitors of sterol biosynthesis. Matching the strong functional conservation of CYP51 activities, the tested imidazoles were found to potently interfere CYP51G1 of Arabidopsis. However, instead of targeting conserved enzymes with C8,7 isomerisation or C24-reductase activities, we found that clomiphene targeted CPI activities in Arabidopsis. Notably, multiple inhibitors of human sterol biosynthesis did not have strong effects on plant growth, indicating that they do not inhibit plant sterol biosynthesis, and were therefore not further analyzed.

Jointly, these data suggests that the exploration of inhibitors of human sterol biosynthesis is a valuable approach for identifying new inhibitors of plan sterol biosynthesis. Exploration of structural variants for improving affinity and specificity could lead to the development of plant-specific sterol biosynthesis inhibitors for agriculture.

### Bifonazole, clotrimazole and econazole are inhibitors of plant CYP51 activity

Multiple azoles were identified as potent fungicides by inhibiting the cytochrome P450-dependent monooxygenase CYP51 (Shafiei *et al*., 2020). They typically act by non-competitive binding to the ferric ion of the heme group of the cytochrome P450, thus preventing substrate binding (Warrilow *et al*., 2013). Sterol C14-demethylation is a highly conserved step in sterol biosynthesis in eukaryotes, and is mediated by a CYP51 homolog. The strong conservation of CYP51 enzymatic activity is also reflected in its sensitivity to different azoles (Shafiei *et al*., 2020). Yet, sequence divergence between fungi, plants and animals has installed differential sensitivities to azoles, allowing their application as fungicides in medicine and agriculture. In example, bifonazole, clotrimazole, and econazole are commonly used to treat fungal infection of the skin and urogenital tracts, reflecting their preference towards fungal over human CYP51s (Warrilow *et al*., 2013). We found that bifonazole, clotrimazole, and econazole interfered with plant growth and development at submicromolar concentrations. Of these inhibitors only clotrimazole was previously shown to be toxic to Lemna species (Alkimin *et al*., 2020), to inhibit radish root growth (Bach, 1985), and root gravitropism in *Pisum sativum* (Amzallag and Vaisman, 2006), without providing evidence that sterol biosynthesis was inhibited in these conditions. Our GC-MS revealed that bifonazole, clotrimazole, and econazole treatments induce the accumulation of sterol biosynthesis intermediates that also accumulate in the Arabidopsis *cyp51* mutant (Kim *et al*., 2005). This provides the first biochemical evidence that these azoles inhibit CYP51 activities in plants. Interestingly, bifonazole and econazole, but not clotrimazole, trigger cell division defects in the root, that are not seen in *cyp51* mutants(Kim *et al*., 2005), suggesting that clotrimazole has a higher specificity towards CYP51.

### Clomiphene is a novel plant sterol biosynthesis inhibitor

Clomiphene is well known as estrogen receptor agonist or antagonist, depending on the target tissue, and is used to induce ovulation or treat breast cancer. In a repurposing screen of FDA approved drugs, clomiphene was identified as an inhibitor of Δ8-7 sterol isomerase and DHCR24 activities (Korade *et al*., 2016). The corresponding enzymes in plants are the Δ8-7 sterol isomerase HYDRA1, and the C24 sterol side chain reductase DWARF1. Consistently with inhibition of sterol biosynthesis enzymes, clomiphene caused altered cell division patterns in the root meristem that were associated with reduced sterol levels. However, the analysis of the sterol biosynthesis intermediates indicated that clomiphene interferes with cyclopropylsterol-cycloisomerase (CPI), and possibly other steps in sterol biosynthesis. Current CPI pharmacology consists of morpholines such as fenpropimorph (Taton *et al*., 1987), and LDAO (Darnet *et al*., 2020). Similarly to clomiphene, neither inhibitor is selective for CPI. Morpholines also inhibit the C14 sterol reductase (FACKLE) and Δ8-7 sterol isomerase (HYDRA1) activities (Taton *et al*., 1987), while LDAO is a potent inhibitor of 2,3-oxidosqualene cyclization to cycloartenol and β-amyrin (Cerutti *et al*., 1985). This suggests that similarity in the biochemical reaction, and in the catalytic center of the different enzymes, make it difficult to develop selective inhibitors. Similarly, clomiphene also caused the accumulation of two other metabolites, indicating that clomiphene also probably has multiple targets in the sterol biosynthesis pathway. In human liver microsomes, 9 clomiphene metabolites could be identified that were more potent estrogen antagonists than clomiphene itself (Mürdter *et al*., 2012). It is therefore not unlikely that clomiphene is also metabolized in plants, and that one or more of these metabolites inhibit one or more steps of plant sterol biosynthesis. A more detailed exploration of clomiphene metabolism in plants will be required. Interestingly, CPI is plant-specific (Desmond and Gribaldo, 2009), indicating that it is an interesting target for developing herbicides.

### A call for caution for using estradiol and SERM-inducible systems

Our analyses identified clomiphene, a Selective Estradiol Receptor Modulator (SERM), as an inhibitor of sterol biosynthesis in plants. Clomiphene caused aberrant cell divisions in the root meristem. Similar cell division phenotypes were observed with tamoxifen, that can be metabolized in human cells into the SERM, 4-hydroxytamoxifen. This suggests that estrogen receptor ligands potentially inhibit sterol biosynthesis in plants.

Several systems that are commonly used for chemically induced expression in plants use estradiol or 4-hydroxytamoxifen as an activating ligand. The most popular system involves the chimeric transcription factor XVE, that can activate a LexA operator based promoter in response to β-estradiol (Zuo *et al*., 2000), that is typically applied between 2 and 10uM in Arabidopsis (Schlucking *et al*., 2013; Wang *et al*., 2020; Yamada *et al*., 2020), and up to 20 uM in rice protoplasts (Chen *et al*., 2017). A derivative hereof allows for activation of UAS operator based promoters in response to the SERM 4-hydroxytamoxifen (Friml *et al*., 2004), that is typically applied at 2uM in Arabidopsis (Kitakura *et al*., 2011). The concentrations of these ligands are in the range of, and even higher than, those at which we observed significant root growth phenotypes for clomiphene and tamoxifen, molecules whose chemical space closely matches that of the inductive ligands, β-estradiol and 4-hydroxytamoxifen, respectively. This indicates that some of the observed phenotypes in such inducible backgrounds may be modified by reduced sterol content, or the accumulation of sterol biosynthesis intermediates. This calls for caution for using such inducible systems in the context of cell biological processes that are sterol-dependent, such as clathrin-mediated endocytosis (Men *et al*., 2008), cytokinesis (Boutte *et al*., 2010; Nakamoto *et al*., 2015) and polarity (Men *et al*., 2008; Stanislas *et al*., 2015)

## Material and methods

### Compounds

All compounds used in the experiments (acitretin, bifonazole, clomiphene, clotrimazole, doxepin, econazole, fluphenazine, flutrimazole, oxiconazole nitrate, perphenazine, tamoxifen, tetraphenylphosphonium, trifluoperazine, trityl chloride) were obtained from Sigma-Aldrich (Overijse, Belgium) and dissolved in DMSO.

### Arabidopsis phenotyping

Gas-sterilized *Arabidopsis thaliana* seeds (Col-0) were plated on ½ Murashige and Skoog (MS) medium supplemented with the appropriate compounds at various concentrations (3 rows/plate, 0.5 cm between seeds). For the primary root length experiments, the plated seeds were first stratified for 3 days in the dark at 4°C and subsequently transferred to a growth chamber under continuous light conditions at 21°C. After 7 days of growth, the plates were scanned and the primary root lengths of the seedlings were measured with Fiji (Schindelin *et al*., 2015). For each treatment, 47-62 individual roots were measured. For the hypocotyl length experiments, the plated seeds were stratified for 3 days in the dark at 4°C. To induce germination they were subjected to 4 hours light, prior to transfer to the dark. After 8 days of growth in the dark, the plates were scanned and the hypocotyl lengths of the seedlings were measured with ImageJ. For each treatment, 33-56 individual hypocotyls were measured.

### GC-MS sterol profiling

*Arabidopsis thaliana* seeds were grown on ½ MS plates for 3 days until germination. Very small seedlings were subsequently transferred to wells of 6-well plates containing 5 ml liquid ½ MS medium and 0.5 or 5 µM of the appropriate compounds, or 0.1% DMSO (1 well per sample, 5 biological replicates per treatment). The seedlings were grown for 5 days in these wells with the compounds, after which they were frozen in liquid nitrogen and thoroughly ground into powder.

Approximately 100 mg (fresh weight) of plant material ground under liquid nitrogen was extracted with 1 mL of methanol to which 5 µg/mL of β-amyrin was added as internal standard. The extractions were carried out at room temperature for 30 minutes, after which the samples were centrifuged at 20,800 x g for 5 minutes. The supernatant was collected and evaporated to dryness under vacuum. The remaining plant material was lyophilized for dry weight determination. The samples were derivatized for GC-MS analysis by adding 10 μL of pyridine and 50 μL of N-Methyl-N-(trimethylsilyl)trifluoroacetamide (Sigma-Aldrich) to the residue. GC-MS analysis was carried out using a GC model 6890 and an MS model 5973 (Agilent). A 1-µL aliquot was injected in splitless mode into a VF-5ms capillary column (Varian CP9013, Agilent). The GC was operated at a constant helium flow of 1 mL per minute and the injector was set to 280°C. The oven was held at 80 °C for 1 minute after injection, then ramped to 280 °C at 20 °C per minute, held at 280 °C for 30 minutes, ramped to 320 °C at 20 °C per minute, held at 320 °C for one minute, and finally cooled to 80 °C at 50 °C per minute. The MS transfer line was set to 250 °C, the MS ion source to 230 °C, and the quadrupole to 150 °C, throughout. Full EI-MS spectra between m/z 60-800 were recorded with a solvent delay of 7.8 minutes. Peak areas were integrated using Masshunter Qualitative Analysis Software (Agilent) and normalized against the dry weight of the sample and the peak area of the internal standard. The total ion currents underneath the peaks corresponding to campesterol, stigmasterol, β-sitosterol, isofucosterol and the internal standard β-amyrin were determined for all samples. To correct for the loss of analyte during sample preparation or analysis, the values for these three sterols were normalized against the internal standard β-amyrin, a triterpene with physicochemical properties similar to the profiled sterols. In addition, the obtained values were also corrected for the amount of plant material that was extracted by dividing the obtained values by the dry weight of the extracted plant material. The relative abundance of the phytosterols β-sitosterol, stigmasterol, campesterol and isofucosterol in the different treatments was calculated by normalization against the DMSO control.

## Acknowledgments

This work was performed in collaboration with the VIB Metabolomics Core and funded by the VIB Innovation Fund, and an FWO research Grant (G001221N). The authors declare no competing interests.

## Notes

### Competing Interest Statement

The authors have declared no competing interest.

